# A novel regulatory pathway recognizes and degrades transcripts with long 3′ untranslated regions

**DOI:** 10.1101/2024.03.11.584429

**Authors:** Ciarán W.P. Daly, Katla Kristjánsdóttir, Jessica D. West, Luyen Tien Vu, Elizabeth A. Fogarty, Rene Geissler, Ciara McDermott, Akila Venkataramany, Andrew Grimson

## Abstract

Quantitative control of gene expression is fundamental to cellular function, and post-transcriptional regulation is a consequential, additional layer of control over protein output. Within mammalian mRNAs, the 3′ untranslated region (3′UTR) acts as a hub of regulatory control, typically mediated by regulatory sequence elements within the 3′UTR. We have found that expression from transcripts with long 3′UTRs is strongly repressed compared to those with short 3′UTRs, and this phenomenon appears to be independent of sequence elements within the 3′UTR. The repression increases with 3′UTR length and is substantial; reporters with 2,000 nucleotide long 3′UTRs are repressed >10-fold compared to 400 nucleotide 3′UTRs. Conversely, increasing the length of the coding region has no effect on expression, demonstrating that it is 3′UTR length, not transcript length, that elicits repression. Reporters with different length 3′UTRs show no difference in transcription rates nor translation efficiency but have clear differences in mRNA half-lives and nucleocytoplasmic distribution, indicating repression of long-3′UTR transcripts is mediated by accelerated RNA decay. However, this transcript degradation does not involve the nonsense-mediated decay (NMD) pathway nor other canonical RNA decay pathways. In order to identify *trans* factors involved in 3′UTR-length-mediated decay, we quantified the differences in proteins bound to reporters with long and short 3′UTRs, and identified over 100 differentially associated factors. Knockdowns of two differentially associating proteins, the mRNA nuclear export factors DDX39B and ZC3H11A, indicate they play a role in repression of mRNAs with long 3′UTRs. This study establishes a novel length-dependent regulatory feature of 3′UTRs, which potentially regulates many genes.

## Introduction

Post-transcriptional regulation of gene expression acts as an additional regulatory layer that enables the fine-tuning of protein output, and allows cells to rapidly respond to changes in their environment (Halbeisen et al. 2008; Mayr 2017). Canonical *trans* factors involved in post-transcriptional regulation act by binding to specific *cis*-regulatory elements within the 3′ untranslated region (3′UTR) of mRNA. In humans and other animals, 3′UTRs contain a diverse set of sequence elements, which recruit regulatory *trans* factors such as microRNAs and RNA-binding proteins (RBPs), which in turn control the stability, localization, and translation efficiency of the mRNA (Mayr 2019). The size and sequence content of over half of human 3′UTRs can be actively regulated through changes in splicing patterns and/or cleavage and polyadenylation, resulting in the generation of alternative 3′UTR isoforms (Lianoglou et al. 2013). These phenomena can alter the fate of the host transcript, changing both the quantitative regulation encoded within a 3′UTR, as well as transcript localization and protein-protein interactions of the encoded protein (Mayr 2019).

Although simpler organisms, such as *Saccharomyces cerevisiae*, usually express genes with relatively short 3′UTRs, in mammals, 3′UTR length approaches the size of the coding sequence, providing extensive sequence space for regulatory elements. For example, human 3′UTRs comprise ∼36% of total mRNA sequence, with an average length of 1,278 nucleotides (nt) (Zhao et al. 2011; Mayr 2017). Notably, over large evolutionary distances, both the length and degree of sequence conservation of 3′UTRs are associated with increased morphological complexity (Chen et al. 2012).

The established paradigm by which 3′UTRs exert their regulatory effects is through the recruitment of functional *trans* factors to *cis*-regulatory sequence elements. However, 3′UTRs may also mediate regulatory outcomes by virtue of their length alone, independent of sequence elements. When analyzed across the transcriptome, increasing 3′UTR length correlates with decreased transcript abundance, although this correlation is inconsistent between different cell types (Yang et al. 2003; Sharova et al. 2009; Spies et al. 2013), perhaps reflecting differences in 3′UTR isoforms and/or alternative complements of *trans* factors. Nevertheless, the underlying cause and mechanism of the association between 3′UTR length and repression is unclear. Recent studies indicate that activating *cis* elements within 3′UTRs are likely as frequent as repressive ones, suggesting that longer 3′UTRs are not repressive merely due to an accumulation of repressive elements (Kristjansdottir et al. 2015; Wissink et al. 2016). The nonsense-mediated decay (NMD) pathway has been shown to target mRNAs with long 3′UTRs (Muhlrad and Parker 1999; Hogg and Goff 2010), but this is contentious (Karousis et al. 2021) and only partially explains the regulatory effects controlled by 3′UTR length.

NMD was initially described as a surveillance mechanism that degrades mRNAs with premature termination codons (PTCs), protecting the cell from producing potentially deleterious polypeptides. However, it is now clear that NMD also acts as a broad regulatory pathway targeting transcripts with a range of triggering features, including normally processed, non-aberrant mRNAs. Aside from a PTC, the two best understood classes of substrates for NMD are transcripts with an intron in the 3′UTR and transcripts with an upstream open reading frame (uORF) (Colombo et al. 2017), both of which are marked by exon-junction complexes (EJCs) bound downstream of terminating ribosomes. The close proximity between a terminating ribosome and an EJC acts as an assembly platform for key NMD factors, ultimately recruiting deadenylases and/or the endonuclease SMG6 (Ohnishi et al. 2003; Eberle et al. 2009; Kervestin and Jacobson 2012). However, NMD can also target mRNAs in an EJC-independent manner (Amrani et al. 2004; Buhler et al. 2006): long 3′UTRs are thought to trigger NMD based on the long physical distance between the terminating ribosome and the poly(A) tail (Behm-Ansmant et al. 2007; Eberle et al. 2008; Ivanov et al. 2008; Silva et al. 2008; Singh et al. 2008). However, compared to canonical NMD, the mechanism behind 3′UTR-length-triggered NMD is poorly understood.

Confounding our understanding of the impact of 3′UTR length on NMD, reporters with artificial 3′UTRs shorter than 1,000 nt long trigger NMD (Eberle et al. 2008; Rebbapragada and Lykke-Andersen 2009; Hogg and Goff 2010), observations that imply a third of human 3′UTRs elicit NMD. However, NMD is estimated to regulate only between 1-10% of human genes (Mendell et al. 2004; Wittmann et al. 2006). One explanation for this dissonance is the presence of “escape” elements (Toma et al. 2015), which, when located at the 5′ end of a 3′UTR, recruit factors that allow NMD substrates to avoid degradation by the NMD machinery (Ge et al. 2016; Kishor et al. 2019; Fritz et al. 2020). However, many transcripts with long 3′UTRs that are not targeted by NMD do not contain identifiable escape elements (Toma et al. 2015), raising the question why only relatively few genes with long 3′UTRs appear to be NMD substrates.

Transcriptome-wide analyses have shown genes with long 3′UTRs are enriched in NMD substrate lists (Bruno et al. 2011; Yepiskoposyan et al. 2011; Hurt et al. 2013). However, recent work using isoform-specific long read sequencing shows no role of 3′UTR length in NMD when discounting transcripts containing an intron in their 3′UTR (Karousis et al. 2021). The conflicting reports on the role of 3′UTR length in NMD make it clear that the influence of 3′UTR length on gene expression is poorly understood, likely due to the challenge of studying 3′UTR length without the confounding impact of *cis*-regulatory elements.

Here, we use random, sequence composition-matched 3′UTRs in reporters to systematically investigate the effect of 3′UTR length on gene expression in human cell lines. We observe substantial repression caused by long 3′UTRs, with a >10-fold difference in expression between reporters with 3′UTRs that differ in length by ∼2,000 nt. We show that this repression results from changes in mRNA stability and demonstrate changes in nucleocytoplasmic distribution of reporters with different length 3′UTRs. Importantly, this repression does not require key NMD factors, but is nonetheless alleviated by escape elements. We identify two *trans* factors involved in 3′UTR length-mediated decay, the mRNA nuclear export factors DDX39B (UAP56) and ZC3H11A. Together, this study supports recent work showing long 3′UTRs alone are not a strong trigger for NMD (Colombo et al. 2017; Karousis et al. 2021), and suggests 3′UTR length acts as global regulatory feature of human genes by modulating mRNA stability independent of NMD.

## Results

### 3′UTR length correlates with decreased expression

The *Hmga2* mRNA harbors an unusually long 3′UTR of ∼3,000 nt, which contains multiple positive and negative regulatory elements (Borrmann et al. 2001; Mayr et al. 2007). While systematically identifying all regulatory sequence elements within the *Hmga2* 3′UTR (Kristjansdottir et al. 2015), we observed that 3′UTR truncations mediated unexpectedly strong repression that appeared unrelated to known repressive *cis*-regulatory elements within the sequence (**Fig 1A**). Indeed, after disrupting all *let-7* miRNA target sites, which constitute the strongest repressive elements within the *Hmga2* 3′UTR, we still observed potent, length-dependent repression (**Fig 1A, red**). These results suggest that the length of this 3′UTR alone, rather than simply the complement of *cis*-regulatory elements within it, is a major determinant of its overall impact on gene expression.

**Fig 1.**
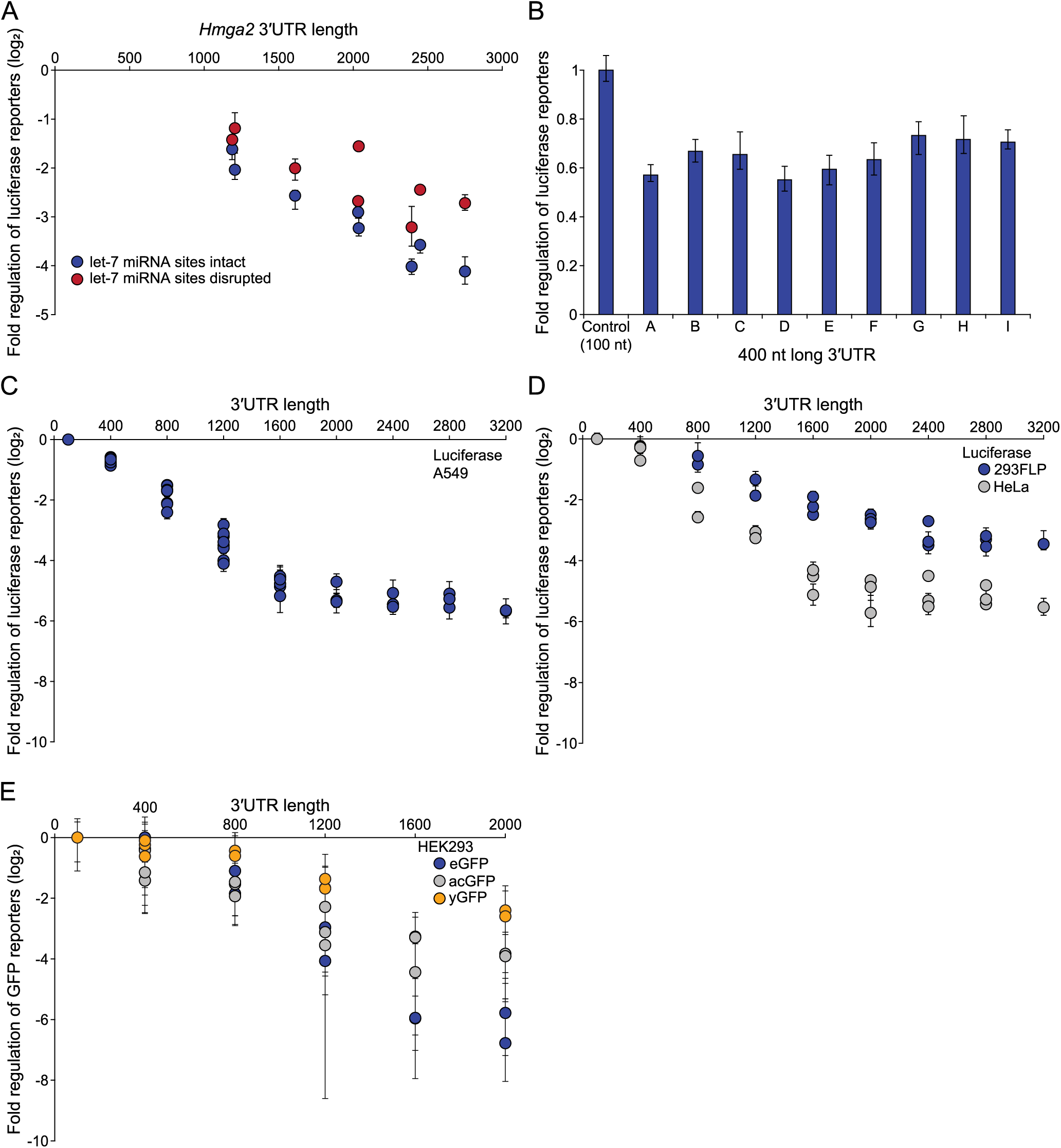
Reporters with long 3′UTRs are repressed in multiple contexts. A) Luciferase assays of *Hmga2* 3′UTR fragments, with and without *let-7* target sites mutated, normalized to three 400 nt random 3′UTRs, in A549 cells (n≥9, error bars show non-parametric estimate of one standard deviation (NP-SD)). B) Luciferase assays of 400 nt random 3′UTRs, normalized to a 100 nt vector-derived control 3′UTR, in A549 cells (n≥9, error bars are NP-SD). C) Luciferase assays of random 3′UTRs concatenated to make progressively longer 3′UTRs, normalized to a 100 nt vector-derived control 3′UTR, in A549 cells (n≥9, error bars are NP-SD). D) As in (C) but in T-REx HEK293FLP (blue) and HeLa (grey) cells. E) Geometric mean fluorescent intensities (gMFIs) of integrated eGFP (blue), acGFP (grey) or yGFP (gold) reporters with different length random 3′UTRs in HEK293 cells, as measured by flow cytometry (>100,000 cells measured, error bars indicate one SD). Each circle represents a unique 3′UTR sequence.

To systematically probe the role of 3′UTR length on gene expression, we designed nine random 400 nt long sequences (400mers) with the same base composition as endogenous human 3′UTRs, but devoid of known RBP binding sites, splice sites, polyadenylation signals and miRNA target sites for highly expressed miRNAs in A549 or HEK293 cells. Our approach was to concatenate these inert 400mers into increasingly longer 3′UTRs, in order to test the effect of 3′UTR length on expression while minimizing the confounding influence of regulatory sequence elements. We first validated that 3′UTR reporters based on each of the nine random 400mers impacted gene expression comparably, as expected for 3′UTRs devoid of functional regulatory elements (**Fig 1B**). Our strategy created a common six nt junction between each concatenated 400mer, which we confirmed had no repressive effect on reporter expression (**Fig S1A**). In total, we generated 52 different 3′UTR sequences, varying in length from 400 to 3,200 nt; a range that encompasses 90% of human 3′UTR lengths. Critically, for each 3′UTR length, we generated multiple 3′UTRs of different sequence, allowing us to better examine the impact of 3′UTR length and not sequence.

We tested the effect of the 3′UTR sequences on expression of luciferase reporters in human A549 cells, and observed a strong negative association between 3′UTR length and expression (**Fig 1C**). This repression was substantial; 2,000 nt long 3′UTRs triggered a ∼20-fold reduction in reporter expression compared to the 400 nt long 3′UTRs. Importantly, reporters with 3′UTRs of the same length mediated similar levels of repression, confirming that length, rather than sequence content, was mediating this correlation. Increasing 3′UTR length for the shorter reporters triggered an approximately linear decrease in expression, which we termed 3′UTR length-mediated decay (LMD). However, reporters containing some of the longest tested 3′UTRs were expressed at similar (albeit very low) levels, suggesting a saturation in the LMD pathway.

To determine if LMD is specific to A549 cells, we tested our 3′UTR reporters in HeLa and HEK293 cells. We observed similar potent repression of reporters with long 3′UTRs, which increased in magnitude with 3′UTR length, although the maximal repression observed varied between cell lines (**Fig 1D**). To rule out potential effects of the coding region, we tested our 3′UTR constructs with three different GFP reporters, including a transcript destabilized, yeast codon-optimized GFP (yGFP). We singly integrated these reporters into the genome using site-specific recombinases at two distinct loci, allowing us to test LMD in a context that mirrors the expression and processing of endogenous mRNAs. Similar to the episomal reporters, these integrated GFP reporters consistently demonstrated substantial 3′UTR length-mediated decay (**Fig 1E**). Thus, reporters with different coding regions, promoters, intron presence/absence and chromatin context, examined across multiple cell lines, persistently demonstrate a repressive effect caused by long 3′UTRs, suggesting a ubiquitous role for 3′UTR length in regulating gene expression in human cells.

### 3′UTR length-mediated decay is not caused by NMD

As nonsense-mediated decay has been shown to target certain mRNAs with long 3′UTRs (Eberle et al. 2008; Singh et al. 2008; Rebbapragada and Lykke-Andersen 2009), we investigated whether NMD was impacting our suite of 3′UTR length reporters. We used multiple different strategies to inhibit NMD, and verified the efficacy of these strategies by measuring the levels of two *SRSF6* splice isoforms, one of which (“PTC+”) is an NMD substrate by virtue of the inclusion of an alternative exon containing a premature termination codon (**Fig S2A, top**). After suppressing NMD using the strategies outlined below, we consistently observed a strong stabilization of the *SRSF6* PTC+ isoform with minimal change in the reference isoform, validating that NMD was effectively inhibited by all strategies employed (**Fig S2A-S2D**).

First, we knocked down *UPF1*, and measured *UPF1* mRNA levels to verify the knockdown (**Fig 2A, inset**). While a luciferase reporter with the *SMG5* 3′UTR (a known NMD target) was stabilized over 2-fold upon *UPF1* knockdown, reporters with different length 3′UTRs showed only relatively minor changes in expression (**Fig 2A**). Importantly, the majority of the repressive effect of LMD was retained, especially in the longer 3′UTR reporters, which retained >95% of the repression observed in control conditions. We observed similar results when knocking down *SMG6* (**Fig 2B**), an additional *trans* factor essential for NMD (Glavan et al. 2006; Eberle et al. 2009).

**Fig 2.**
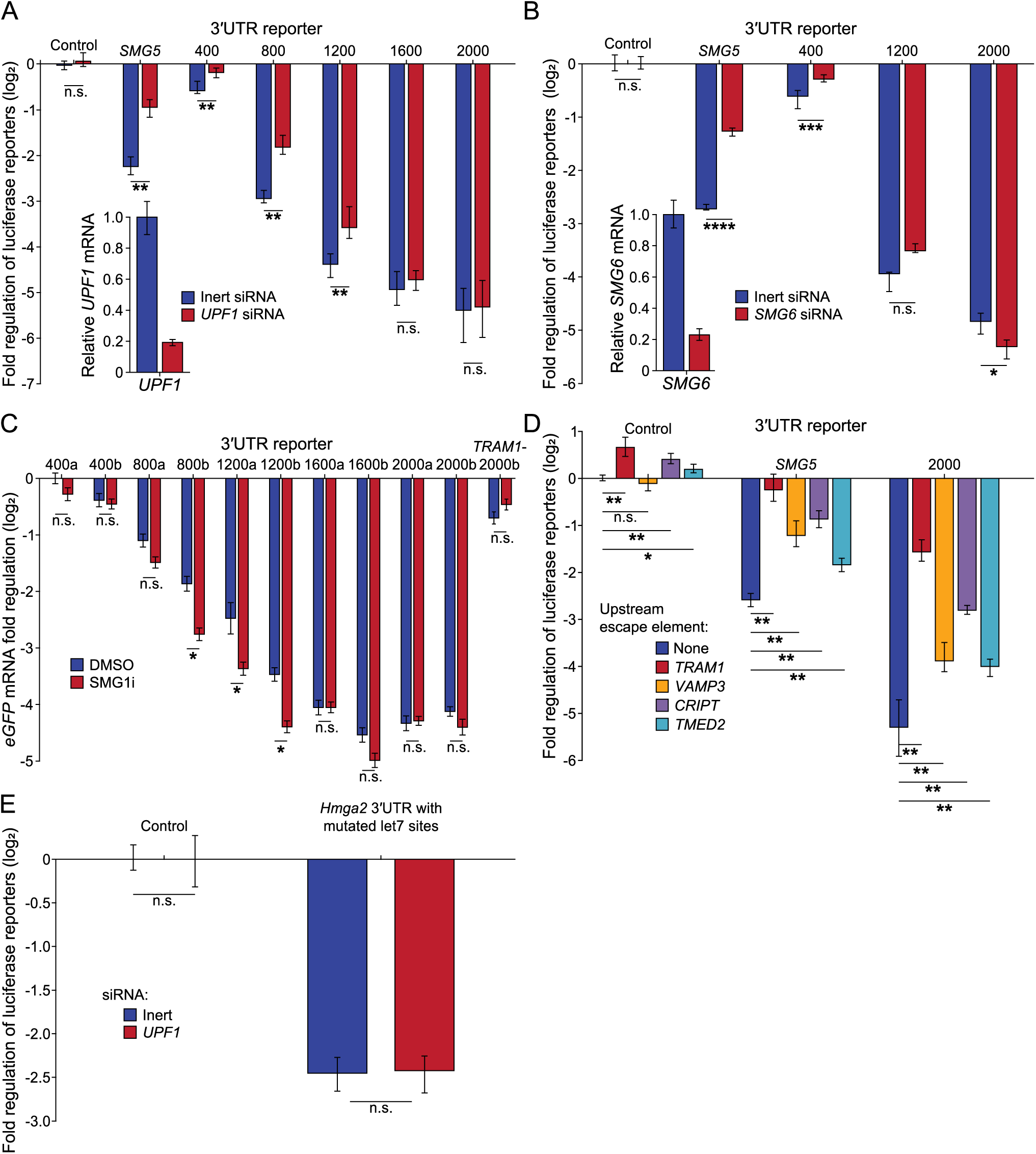
NMD does not cause repression of long 3′UTR reporters. A) The effect of UPF1 knockdown on expression of transiently transfected luciferase reporters with the *SMG5* 3′UTR and different length random 3′UTRs, normalized to a 100 nt vector-derived control 3′UTR (n≥9, three independent siRNA transfections, error bars are NP-SD). Inset: Relative *UPF1* mRNA levels after 72-hour siRNA treatment in HeLa cells, compared to inert siRNA, as measured by RT-qPCR (n=9, error bars indicate one SD). B) As in (A) but upon treatment with *SMG6* siRNA and showing *SMG6* mRNA levels in inset. C) The effect of NMD inhibition by a four hour incubation with 0.5 µM SMG1i on mRNA levels of integrated eGFP reporters with different length random 3′UTRs, including a 2,000 nt 3′UTR with an upstream *TRAM1* escape element, in HEK293T cells, relative to DMSO vehicle treatment, as measured by RT-qPCR (three biological replicates each in technical triplicate, error bars are one SD). “a” and “b” signify random 3′UTRs of the same length but different sequence. Normalized to the 400a random 3′UTR reporter with DMSO treatment. D) The effect of incorporating NMD escape elements from indicated genes (see legend) to the 5′ end of the 3′UTR of luciferase reporters with a 100 nt, 2,000 nt random, or *SMG5* 3′UTR in A549 cells (n=9, error bars are one NP-SD). Normalized to the 100 nt vector-derived control 3′UTR with no upstream escape element. E) The effect of *UPF1* knockdown on expression of transiently transfected luciferase reporters expressing the *Hmga2* 3′UTR with mutated *let-7* target sites, normalized to a 100 nt vector-derived control 3′UTR, in HeLa cells (n≥9, three independent siRNA transfections, error bars are one NP-SD). *p<0.05, **p<0.01, ***p<0.001, ****p<0.0001; p-values derived from two-tailed t-tests.

UPF1 and SMG6 have roles beyond NMD (Kim and Maquat 2019), so knocking them down will likely have consequences not limited to suppression of NMD. Therefore, we also treated cells with a small molecule, SMG1i, designed to specifically inhibit NMD. SMG1i acts by blocking the kinase activity of SMG1 (Gopalsamy 2012, Mino 2019), which is required for NMD (Langer et al. 2021). Consistent with our previous results, SMG1i treatment stabilized the “PTC+” isoform of *SRSF6* (**Fig S2D**) but had minimal effect on the expression of different length 3′UTR reporters (**Fig 2C**). Altogether, these results show that NMD plays little to no role in the repressive action mediated by these long 3′UTR reporters, in contradiction to previous studies.

Our final approach to explore the connection between NMD and LMD was to test NMD escape elements, which are approximately 200 nt long and located at the 5′ end of the 3′UTR. Escape elements permit NMD substrates to avoid degradation (Toma et al. 2015), likely through the recruitment of NMD antagonistic factors such as PTBP1 and hnRNP-L (Ge et al. 2016; Kishor et al. 2019). Surprisingly, given our NMD knockdown results, we observed a strong stabilization of reporters with 2,000 nt long 3′UTRs when different escape elements were introduced downstream of the reporter stop codon (**Fig 2D**). As is true for NMD substrates (Toma et al. 2015), these escape elements were only effective when located at the start of the 3′UTR (**Fig S2E**). However, their protective effect was not dependent on UPF1 (**Fig S2F**). The substantial NMD-independent stabilization of long 3′UTRs by escape elements suggests that escape elements are not specific to NMD.

Finally, to determine whether length-mediated repression by the full-length *Hmga2* 3′UTR was sensitive to NMD, we examined the effect of UPF1 siRNA knockdown on expression of the *Hmga2* 3′UTR reporter (**Fig 2E**). For these experiments, we used the *Hmga2* 3′UTR reporter with mutated *let-7* target sites to focus the assay on testing the repressive effect of 3′UTR length specifically, without being confounded by the substantial repression mediated by these microRNA target sites. Importantly, loss of UPF1 had no impact on the expression of the Hmga2 3′UTR reporter, demonstrating that NMD is not required for the strong repressive activity mediated by this long 3′UTR.

### LMD involves accelerated mRNA decay

After establishing that NMD is not the primary pathway through which long 3′UTR reporters are repressed, we set out to understand how long 3′UTRs trigger LMD. We first asked whether the length of the 3′UTR or the length of the entire transcript triggers LMD. We compared expression of reporters of similar transcript length while varying either the 3′UTR length or the length of the coding region (CDS). We observed that changing the length of the CDS had little effect on reporter expression compared to changing the 3′UTR length (**Fig 3A**), suggesting that LMD senses 3′UTR length rather than total transcript length.

**Fig 3.**
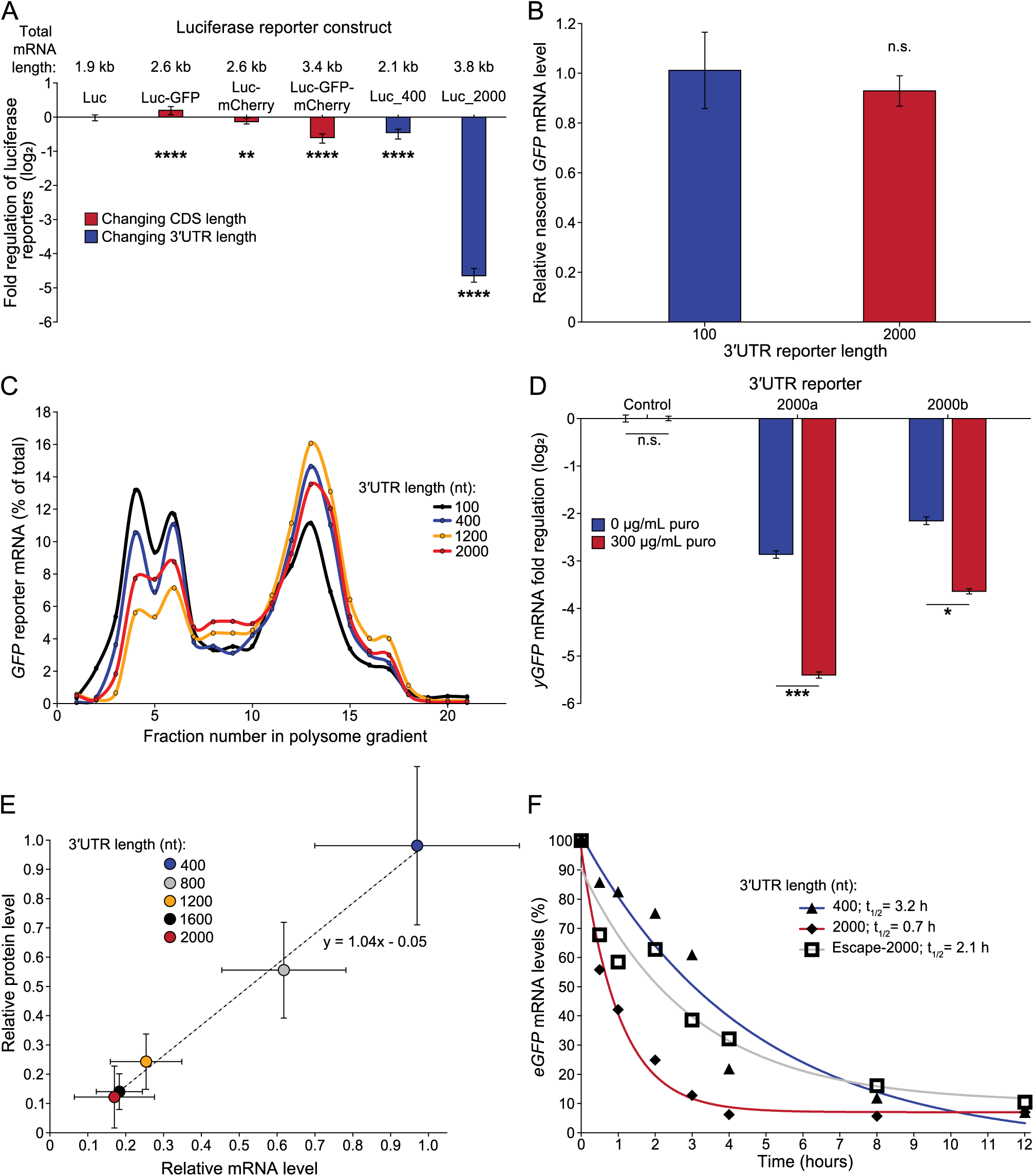
LMD destabilizes the mRNA of reporters with long 3′UTRs. A) Luciferase assays of reporters with different length coding sequences or random 3′UTRs in A549 cells. CDS length of luciferase reporters was increased by downstream incorporation of the CDS from fluorescent proteins, separated by 2A peptides, and all CDS length constructs contain a 400 nt random 3′UTR. Normalized to luciferase reporter with a 100 nt vector-derived 3′UTR (n=18, error bars are one NP-SD). Statistical tests compare Luc sample against indicated sample. B) Relative nascent mRNA levels of integrated GFP reporters with different length random 3′UTRs as measured by RT-qPCR of mRNA isolated through transcriptional run-on reactions with biotin NTPs (see Methods) in T-REx HEK293FLP cells (two biological replicates each in technical triplicate, error bars are one SD). C) Polysome profiles of integrated GFP reporters with different length random 3′UTRs in T-REx HEK293FLP cells (one biological replicate in technical triplicate). D) The effect of translation inhibition by an 8 hour treatment with 300 µg/mL puromycin on the mRNA levels of two integrated yGFP reporters with distinct 2,000 nt random 3′UTRs in T-REx HEK293FLP cells, as measured by RT-qPCR (three biological replicates each in technical triplicate, error bars are one SD). E) Comparison of relative protein and mRNA levels of integrated GFP reporters with different length random 3′UTRs in T-REx HEK293FLP cells, normalized to the average of three 400 nt random 3′UTRs. mRNA levels measured by RT-qPCR (three biological replicates each in technical triplicate, error bars indicate one SD), protein levels measured by flow cytometry (>100,000 cells measured, error bars indicate one SD). At least three different random 3′UTR reporters were used per indicated length. F) mRNA decay curves of integrated eGFP reporters with different length random 3′UTRs expressed from the doxycycline repressible Tet-OFF promoter in HEK293T cells, measured by RT-qPCR (three biological replicates each in technical triplicate). Half-lives of each reporter noted in legend. *p<0.05, **p<0.01, ***p<0.001, ****p<0.0001; p-values derived from two-tailed t-tests.

To understand which stage in the mRNA lifecycle is acted upon by LMD, we tested whether transcription of 3′UTR reporters of different lengths is consistent. We performed nuclear run-on reactions in the presence of biotinylated NTPs, then isolated and quantified nascent transcripts from actively transcribing polymerase through the biotin molecular handle (Kwak et al. 2013). We observed little difference in the levels of nascent mRNA of reporters with a long 2,000 nt 3′UTR compared to a short 3′UTR (**Fig 3B**), indicating the reporters are transcribed at similar efficiencies regardless of 3′UTR length.

To test whether reporters with long 3′UTRs have lower expression because of changes in translation, we stably expressed our reporters in HEK293 cells and measured their translation efficiency with polysome profiling. We observed similar polysome profiles for reporters with 3′UTRs that varied in length from 100 to 2,000 nt (**Fig 3C**), demonstrating no major difference in translation efficiency as a function of 3′UTR length. As an alternative approach, we inhibited translation with either puromycin (**Fig 3D**) or cycloheximide (**Fig S3A**), two small molecules that block translation through different mechanisms of action (Pestka 1971; Schneider-Poetsch et al. 2010). If the repressive function of LMD was acting through translation, we would expect that inhibition of translation would stabilize reporters with long 3′UTRs. Interestingly, treatment with either drug substantially downregulated two distinct long (2,000 nt) 3′UTR reporters (**Fig 3D & S3A**). These data suggest translation is not necessary for LMD, and additionally implies that translating mRNPs may be somewhat protected from LMD.

To confirm that the repression of reporters with increasingly longer 3′UTRs is not occurring through changes in translation, we compared the relative mRNA and protein levels of our reporters. We observed a strong linear correlation between mRNA and protein levels with a slope of ∼1 (**Fig 3E**), suggesting that mRNA levels alone dictate protein levels of the 3′UTR reporters. Together, these results demonstrate that lengthening the 3′UTR in reporters has no effect on translation nor transcription, implying that LMD acts by altering the levels of mature mRNA.

To test the hypothesis that LMD destabilizes mature mRNAs with long 3′UTRs, we measured the mRNA half-lives of reporters with different length 3′UTRs. We used the repressible Tet-Off promoter to block reporter transcription with doxycycline, and used a pulse-chase approach to monitor mRNA decay. We observed a striking difference in reporter half-life (**Fig 3F & S3B**), with a ∼3 hour half-life for a 400 nt 3′UTR reporter compared to just ∼1 hour for a 2,000 nt long 3′UTR. Furthermore, incorporating an escape element upstream of a 2,000 nt long 3′UTR approximately doubled the reporter half-life (**Fig 3F**). Thus, the LMD pathway results in the destabilization of mRNAs with long 3′UTRs.

### Nuclear export factors are required for LMD

To identify factors involved in driving the destabilization of reporters with long 3′UTRs, we used validated shRNAs (Geissler et al. 2016) to knock down two major exonucleases (*XRN1* and the exosome), as well as the deadenylases *CNOT1*, *PAN3* and *PARN*, but did not observe any substantial stabilization of long 3′UTR reporters relative to short 3′UTR reporters (**Fig S3C**). Thus, none of the established, major RNA decay pathways seem to contribute to LMD.

Given that canonical RNA decay pathways do not appear to contribute to LMD (**Fig S3C**), we took a biochemical approach to identify factors that might underlie the destabilization of reporters with long 3′UTRs. Comprehensive Identification of RNA-binding Proteins by Mass Spectrometry (ChIRP-MS) leverages antisense biotinylated probes tiled against a transcript of interest to identify associated proteins (Chu et al. 2015; Chu and Chang 2018). To quantitatively compare the association of proteins associated with *GFP* reporters with a 400 nt 3′UTR, 2,000 nt 3′UTR and 2,000 nt long 3′UTR with an upstream escape element, we used Stable Isotope Labelling by Amino acids in Cell culture (SILAC). SILAC-labelled HEK293 cells were transfected, pooled, *GFP* mRNPs were isolated by ChIRP, and then associated proteins were identified and deconvoluted by mass spectrometry (**Fig 4A**). Our rationale was that factors involved in LMD would associate differentially with each of the three different reporters. Across two biological replicates, ChIRP isolated approximately one third of *GFP* mRNA in the pooled sample (**Fig S4A**), and the proteins identified by mass spectrometry were strongly enriched for factors involved in RNA biology (**Fig S4B**), suggesting that SILAC-ChIRP-MS faithfully identified proteins bound to mRNA of the GFP reporters. We verified that the long 3′UTR was not being spliced into a shorter 3′UTR (**Fig S3D**). Over 200 proteins were associated with each reporter mRNA (identified by >1 peptide), in line with previous studies of mRNPs (Muller and Glaunsinger 2017).

**Fig 4.**
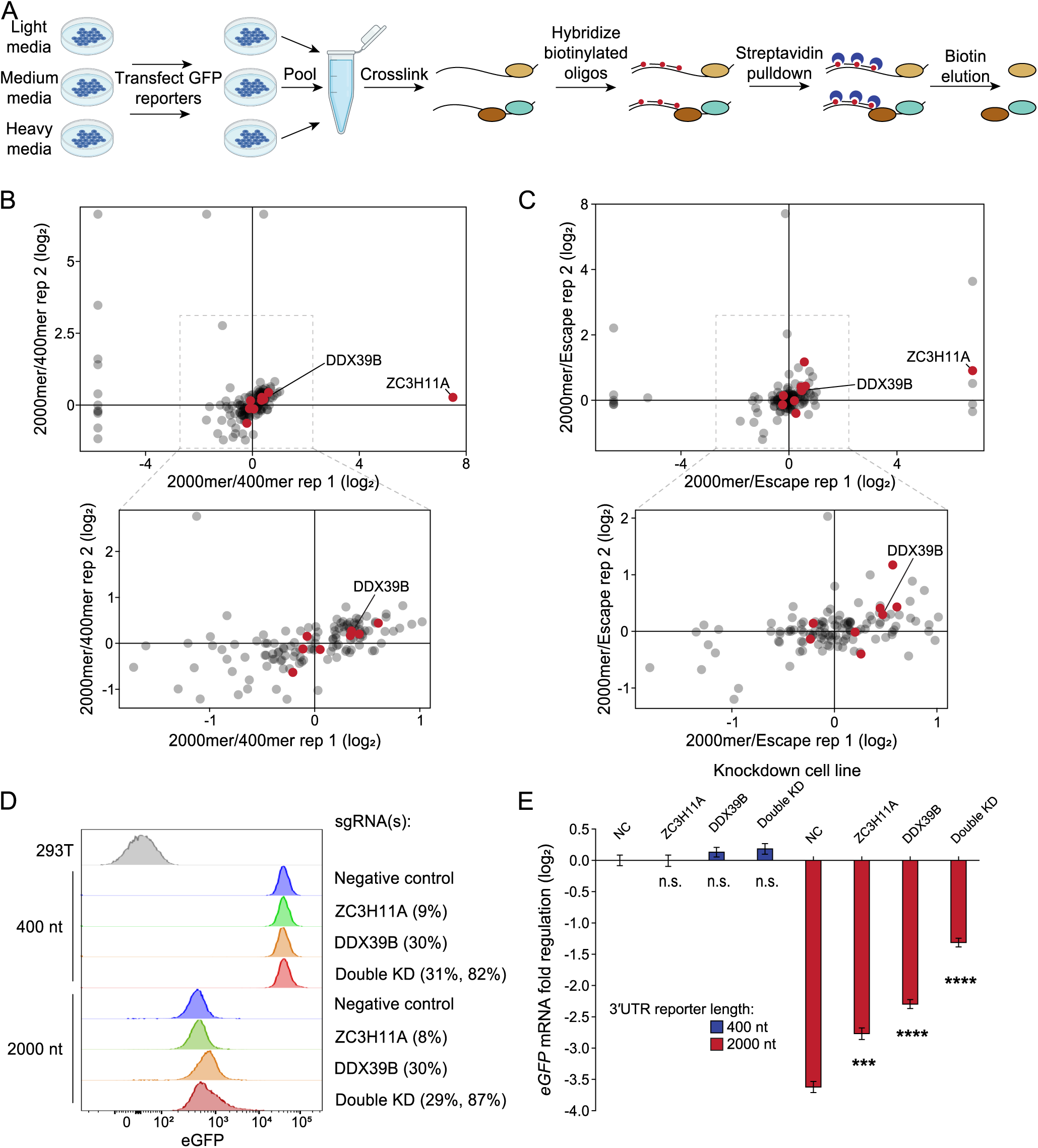
SILAC-ChIRP-MS identifies DDX39B and ZC3H11A enrichment on long 3′UTR reporter. A) Experimental outline of SILAC-ChIRP-MS in T-REx HEK293FLP cells transiently transfected with GFP reporters with either a 400 nt random 3′UTR, 2,000 nt random 3′UTR or 2,000 nt random 3′UTR with an upstream *TRAM1* escape element. B) SILAC abundance ratios of proteins associated with the GFP reporter with a 2,000 nt random 3′UTR relative to the GFP reporter with a 400 nt random 3′UTR, comparing two SILAC-ChIRP-MS replicates. Bottom shows zoomed view of boxed area. Red dots indicate proteins part of the gene ontology term “mRNA export from nucleus”. C) As in (B), but ratios of proteins associated with GFP reporter with a 2,000 nt random 3′UTR relative to GFP reporter with 2,000 nt random 3′UTR with an upstream *TRAM1* escape element. D) Distribution of eGFP fluorescence intensities as measured by flow cytometry (>100,000 cells measured), for HEK293T cells expressing integrated eGFP reporters with 400 nt or 2,000 nt random 3′UTRs under labelled CRISPRi knockdown conditions. Percentages indicate knockdown levels as measured by RT-qPCR (three biological replicates, each measured in technical triplicate), with the double KD showing ZC3H11A and DDX39B co-knockdown levels. E) Reporter mRNA levels in HEK293T cells expressing an integrated eGFP reporter with 400 nt or 2,000 nt random 3′UTRs under labelled CRISPRi knockdown conditions (knockdown levels shown in Fig 4D), as measured by RT-qPCR (three biological replicates in technical triplicate, error bars are one SD). Statistical tests compare cell line expressing a negative control sgRNA (NC) against indicated sample for each reporter. ***p<0.001, ****p<0.0001; p-values derived from two-tailed t-tests.

We compared the enrichment of proteins associated with the long 3′UTR reporter against those associated with the short 3′UTR reporter (**Fig 4B**) or the long 3′UTR reporter with an upstream *TRAM1* escape element (**Fig 4C & S4C-D**). Proteins enriched on the long 3′UTR reporter were ranked by their SILAC fold changes, and rankings across the two ChIRP-MS replicates were combined to build a list of differentially associated factors. We reasoned that factors involved in LMD would be consistently enriched, or consistently depleted, when comparing the long 3′UTR reporter against both the short 3′UTR and the long 3′UTR with an escape element. Thus, the list of putative *trans* factors involved in LMD was further refined following this rationale, by aggregating the ranked scores of the most enriched or depleted factors across the two comparisons. Based on this list, we screened approximately 70 factors for their role in LMD through CRISPR interference (CRISPRi) knockdowns, and measured the effect on reporters with short and long 3′UTRs (**Fig S4E**).

Knockdown of DDX39B (UAP56), a protein with known roles in mRNA nuclear export and splicing (Luo et al. 2001), increased expression of eGFP protein from reporters with a long, but not a short, 3′UTR (**Fig 4D**). Furthermore, knockdown of DDX39B led to a ∼3-fold stabilization of *eGFP* mRNA from the reporter with a 2,000 nt random 3′UTR, with no change in the mRNA levels of the 400 nt random 3′UTR reporter (**Fig 4E**). We noticed a slight enrichment of factors involved in mRNA export associated with the long 3′UTR reporter (**Fig 4B-C, red dots**), with a particularly strong enrichment of ZC3H11A, a component of the nuclear export complex TREX (Folco et al. 2012). Concurrently knocking down both DDX39B and ZC3H11A upregulated expression of eGFP from the reporter with a long 3′UTR (**Fig 4D**), and led to a ∼4-fold stabilization of the *eGFP* mRNA (**Fig 4E**). Therefore, both DDX39B and ZC3H11A, components of the same nuclear export complex, appear to play a role in regulating expression from transcripts with long 3′UTRs.

Given the discovery of TREX components as *trans* factors involved in LMD, we used RNA FISH to measure reporter mRNA levels in the nuclear and cytoplasmic compartments. We observed significant differences in the nuclear/cytoplasmic distribution of reporter mRNA with different length 3′UTRs (**Fig 5A-C**), where transcripts with longer 3′UTRs were increasingly nuclear; ∼35% of transcripts for a 400 nt long 3′UTR reporter localized to the nucleus compared to ∼55% nuclear for a 2,000 nt long 3′UTR (**Fig 5B**). Control FISH probes against *GAPDH* showed minimal difference in the cellular distribution of *GAPDH* mRNA across reporter cell lines, as expected (**Fig S5A-B**). Interestingly, the RNA FISH images also show nuclear puncta present only in cells expressing reporters with long, but not short, 3′UTRs (**Fig 5C, bottom**). Since these puncta were restricted to a single punctum always in the nucleus, they may correspond to the site of transcription for these reporters, which are integrated and hemizygous, although we cannot rule out transcript aggregation in phase-separated bodies. Overall, our results indicate that LMD decreases the half-life of reporters with long 3′UTRs, and suggest 3′UTR length alters mRNA distribution in the cell, potentially through changes in nuclear export efficiency mediated by the TREX complex.

**Fig 5.**
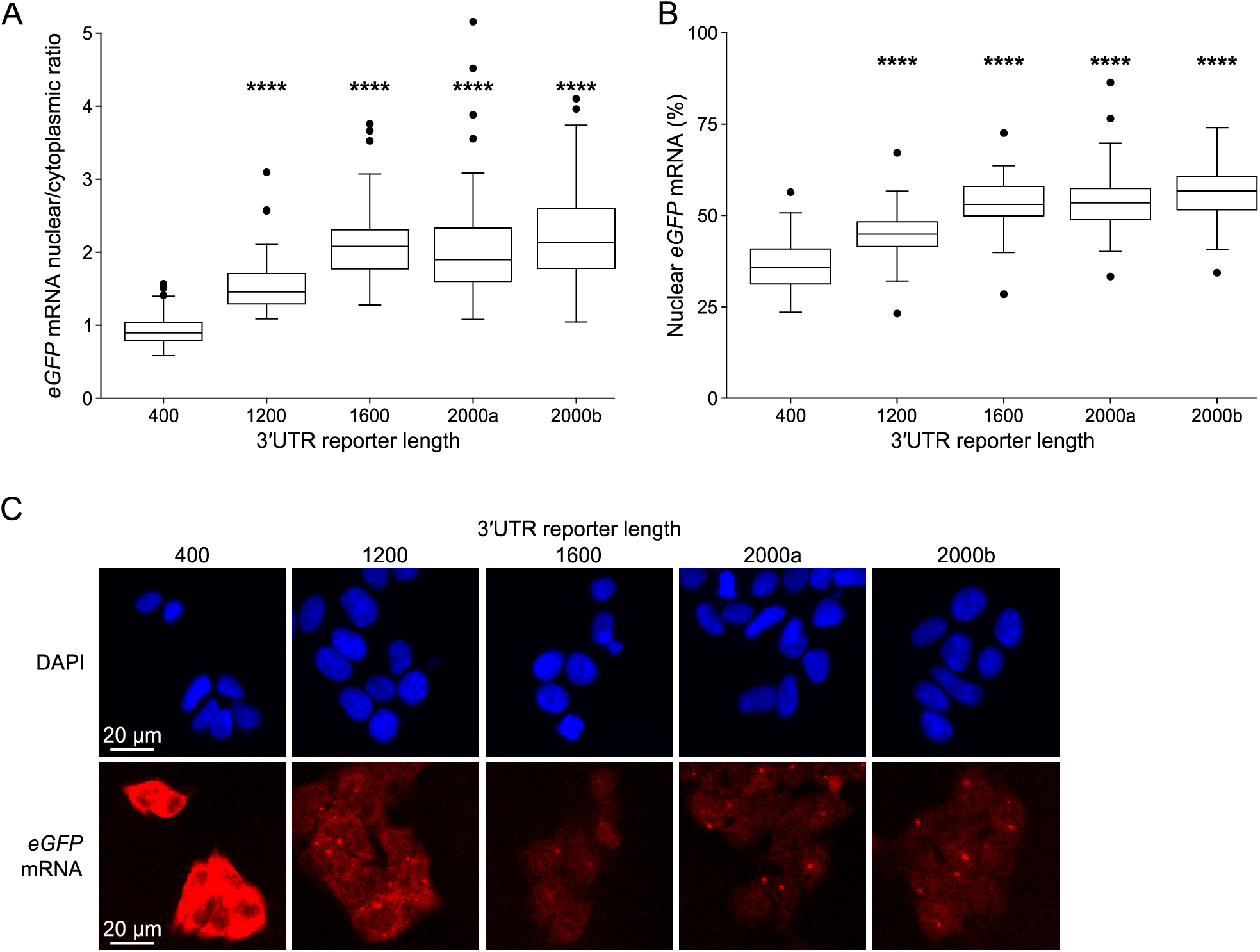
3′UTR length alters nucleocytoplasmic distribution. A) Nuclear/cytoplasmic ratios of reporter mRNA in HEK293T cells expressing an integrated eGFP reporter with different length random 3′UTRs, as measured by RNA FISH (n≥50 cells). Statistical tests compare 400 nt long 3′UTR reporter against indicated reporter. B) The percent of *eGFP* mRNA localized to the nucleus for different length random 3′UTR reporters, as measured by RNA FISH (n≥50 cells). Statistical tests compare 400 nt long 3′UTR reporter against indicated reporter. C) Representative images of DAPI stained (top) and RNA FISH labelled (bottom) HEK293T cells expressing integrated eGFP reporters with different length random 3′UTRs. ****p<0.0001; p-values derived from two-tailed t-tests.

## Discussion

Human 3′UTRs range in size over three orders of magnitude (Zhao et al. 2011; Mayr 2017), and their length is frequently regulated through APA. More than half of human genes are estimated to express alternative 3′UTRs (Lianoglou et al. 2013), which drive regulatory changes across cell types and in different environmental contexts. Considering the median human 3′UTR size is ∼1,200 nt, any regulatory mechanism targeting long 3′UTRs has the potential to impact a substantial portion of the human transcriptome. Nevertheless, the extent to which 3′UTR length contributes to gene regulation has long been unclear. Previous studies have demonstrated a trend towards decreased stability for transcripts with long 3′UTRs in some cell lines (Yang et al. 2003; Sandberg et al. 2008; Spies et al. 2013) but not others (Sharova et al. 2009). The destabilization of long 3′UTRs has historically been attributed to an increased number of repressive elements or to the triggering of NMD. Here, we establish that length is a regulatory feature inherent to 3′UTRs, with progressively longer 3′UTRs eliciting increasingly potent decay. This trend was consistent across cell lines and gene context, and, importantly, independent of NMD.

Our strategy relied upon random sequences to design artificial 3′UTRs of varying lengths that were devoid of *cis*-regulatory elements and contained the same GC content as endogenous human 3′UTRs. This strategy allowed us to study 3′UTR length uncoupled from the impact of *cis*-regulatory effects that confound other studies. Previous studies often investigated 3′UTR length by inverting endogenous 3′UTR sequences (Schmidt et al. 2015) or by the addition of unrelated sequences to the 3′UTR (Chatterjee and Pal 2009; Popp and Maquat 2013), without assessing for the presence of regulatory motifs when creating these artificial 3′UTRs. Across our experiments, we used multiple 3′UTRs of identical length but different sequence content to ensure any observed regulatory effects were specifically driven by 3′UTR length. Using this approach, we demonstrate that 3′UTR length-mediated decay alters half-life and nucleocytoplasmic distribution of the corresponding transcripts.

We tested whether NMD was driving 3′UTR length-mediated decay of our reporters by multiple distinct approaches, all of which confirmed this was not the case. This is in sharp contrast to previous studies investigating the impact of 3′UTR length on NMD using reporters, which showed that artificial 3′UTRs >800 nt trigger NMD (Buhler et al. 2006; Eberle et al. 2008; Hogg and Goff 2010; Yepiskoposyan et al. 2011). A recent study robustly identified NMD targets through combined knockdowns and rescue of UPF1, SMG6 and SMG7 followed by both short- and long-read sequencing (Colombo et al. 2017; Karousis et al. 2021). This study observed no correlation between 3′UTR length and NMD after accounting for long 3′UTRs containing introns. Karousis and colleagues highlighted that the inclusion of long-read sequencing was critical for accurately defining 3′UTR length, since typical annotations would overlook the widespread alternative 3′UTR isoform usage caused by knockdown conditions. This is especially relevant considering many splicing factors are themselves targets of NMD (García-Moreno and Romão 2020). Although this study disputes previous transcriptome-wide identification of NMD substrates by 3′UTR length, the breadth of reporter data showing that NMD targets long 3′UTRs remains credible. While it is challenging to explain the discrepancy in 3′UTR length reporter studies, recent advances in the understanding of NMD have made it clear that NMD is not a single pathway but instead an interconnected network of decay pathways linked by a shared set of *trans* factors (Karousis and Muhlemann 2022). It is possible that LMD could be associated with this network in some way, albeit without the involvement of UPF1, SMG1 or SMG6, three core NMD factors. Moreover, we suspect that previous reporter studies that attributed repression mediated by long 3′UTRs to NMD may, in fact, have been detecting LMD, or perhaps a combination of LMD and NMD.

Both our microscopy data and the role of two nuclear export factors in LMD imply long 3′UTRs trigger decay at some point during or shortly after nuclear export. Current mRNP composition models make it difficult to explain how 3′UTR length could be sensed in the nucleus without actively translating ribosomes marking the stop codon, so we hypothesize that LMD occurs soon after nuclear export, potentially before the first completed round of translation in the cytoplasm. Translation appears to have a protective effect from LMD, as shown by our experiments using drugs to inhibit translation, which unexpectedly increased the potency of LMD. Therefore, the small fraction of transcripts with long 3′UTRs that, perhaps stochastically, escape LMD and enter the pool of translating mRNAs may then be subsequently protected from LMD, as has been shown for subpopulations of NMD substrates (Hoek et al. 2019). While it is clear that LMD reduces the half-lives of reporters with long 3′UTRs (**Fig 3F**), the underlying mechanism responsible for this destabilization is yet to be explained. DDX39B and ZC3H11A are both components of the TREX complex, and a recent structural study has shown how TREX and EJC components multimerize to compact mRNPs for nuclear export (Pacheco-Fiallos et al. 2023). We propose a model whereby long 3′UTRs, devoid of EJCs, are poorly packaged before nuclear export, leading to the retainment of certain nuclear export factors which then elicit decay soon after nuclear export.

The distribution of 3′UTR lengths is known to exhibit marked tissue-specificity, with an unusual abundance of transcripts with long 3′UTRs found in neurons (Wang and Yi 2014) and a depletion of mRNAs with long 3′UTRs in spermatocytes (Liu et al. 2007). In neurons, long 3′UTRs have been shown to trigger inflammation by recognition of dsRNA by antiviral response pathways, likely due to increased RNA secondary structure in long 3′UTRs (Dorrity et al. 2023). In contrast, 3′UTR shortening in spermatocytes is caused by changes in APA (Liu et al. 2007) and a poorly understood non-canonical NMD pathway (Bao et al. 2016). Furthermore, global shortening of 3′UTR length has been shown to occur in numerous cancer types (Mayr and Bartel 2009; Fan et al. 2020; Yu et al. 2022; Chan et al. 2023). Studying 3′UTR length has inherent value due to its potential to regulate a considerable portion of the transcriptome, but these examples also demonstrate the impact of 3′UTR length in specific tissues and in disease. Further work is needed to disentangle LMD and NMD, and to understand how the cell identifies long 3′UTRs and marks them for degradation.

## Methods

### Cloning and Plasmids

#### Random 400mer 3′UTR luciferase reporters

The sequences of the random 400mer 3′UTR mimics were generated to mimic the base composition of human 3′UTRs (A:27%, U:29%, C:22%, G:22%), excluding sequences containing target sites for the top 20 expressed miRNAs in the cell lines used. Nine random 400mer sequences were chosen and synthesized as gBlocks (IDT), with SpeI and NheI-NotI restriction sites (5′ and 3′, respectively) of the random sequence. Each 400mer was inserted downstream of the firefly luciferase coding sequence and upstream of the SV40 late poly(A) signal of Promega’s pmirGLO Dual-Luciferase miRNA Target Expression vector, using SpeI and NotI digested fragments ligated into the the NheI and NotI sites within the vector.

#### Generating larger random sequence 3′UTR reporters

During the cloning of the random 400mer 3′UTR reporters, the original vector NheI site is destroyed. However, the insert adds a new NheI site downstream of the random 400mer sequence and immediately upstream of the NotI site. This allows for sequential cloning of random 400mers using the same strategy. Random 400mer 3′UTR mimic plasmids are digested with NheI and NotI and another random 400mer insert, digested with SpeI and NotI, is cloned downstream of inserts cloned previously, again destroying the upstream NheI site and adding a new one downstream.

#### Random 3′UTR acGFP reporters

Random 400mers and their concatamers were isolated from luciferase reporters using PCR, replacing the restriction sites with NotI and PmeI. Inserts were then cloned between NotI and PmeI restriction sites in an acGFP/dsRed pEF5/FRT/V5-D-TOPO derived plasmid (Wissink et al. 2016).

#### yGFP reporters

The GFP open reading frame in each random 3′UTR mimic reporter was replaced with a yeast codon-optimized GFP, PCR amplified from the pFA6a-GFP(S65T)-Trp1 plasmid, a kind gift from John Pringle (Addgene plasmid# 41597) (Choe et al. 2011). A start codon was added to the pFA6a-GFP-Trp1-derived GFP during PCR amplification.

#### eGFP reporters

Random 400mers and their concatemers were subcloned by Gibson assembly downstream of eGFP in the attB-containing pFL2 vector, a kind gift from Douglas Fowler.

#### NMD escape sequences

The first 200 nts of the human TRAM1 and VAMP3 3′UTRs were isolated from genomic DNA using nested PCR, adding a PmeI site upstream and NheI and NotI sites downstream. Amplified inserts were cloned into pmirGLO using PmeI and NotI sites. The resulting plasmid was digested with NheI and NotI to add the SMG5 3′UTR or random sequence 3′UTRs downstream. The SMG5 3′UTR was cloned from genomic DNA using nested PCR, adding a SpeI restriction site upstream and NheI and NotI sites downstream.

#### Reference 3′UTR control reporter

Control reporters were generated by digesting the pmirGLO vector with NheI and NotI, or the GFP vector with NotI and PmeI, and using T4 DNA polymerase to create blunt ends, which were then ligated. The resulting luciferase and GFP reporters include ∼175 nts and ∼206 nts of 3′UTR sequence, respectively, which are also present in all derivative constructs.

#### CDS length reporters

CDS length was increased by cloning fluorophore coding sequences downstream of firefly luciferase in Promega’s pmirGLO Dual-Luciferase miRNA Target Expression vector, separating the coding regions by P2A or T2A sequences ordered from IDT. GFP CDS was subcloned from the above GFP vectors, mCherry CDS was subcloned from pHR-UCOE-SFFV-dCas9-mCherry-ZIM3-KRAB, a kind gift from Mikko Taipale (Addgene plasmid #154473) (Alerasool et al. 2020). Cloning was performed with Gibson assembly.

#### sgRNA vectors for CRISPRi

sgRNA sequences, including negative control non-targeting sequences, were taken from the hCRISPRiv2 library (Horlbeck et al. 2016) and ordered as oligos from IDT. For single sgRNA expression, the sgRNA oligos were cloned by Gibson assembly into the pU6-sgRNA EF1a-puro-T2A-BFP vector, a kind gift from Jonathan Weissman (Addgene #60955) (Gilbert et al. 2014). For co-expression of sgRNAs, the sgRNA oligos were cloned by Gibson assembly into a homemade version of the three-guide Perturb-seq vector designed by Britt Adamson and colleagues (Adamson et al. 2016), a kind gift from Divya Ganapathi Sankaran.

All plasmids were sequence verified by either Sanger sequencing or whole-plasmid sequencing provided by Plasmidsaurus (Oregon, USA). All plasmids are available upon request.

### Luciferase assays

Cells grown in 24-well plates in DMEM supplemented with 10% FBS (Gibco) were seeded 24 hours before transfection, at the following densities: 75,000 cells/well for A549 cells, 150,000 cells/well for Flp-In^TM^ T-REx^TM^ HEK293 cells and 50,000 cells/well for HeLa cells. Plasmids (equimolar amounts, 14-20 ng/well depending on plasmid size) were transfected into cells, along with carrier DNA to adjust the total amount of DNA (100-140 ng/well pUC19), using Lipofectamine 2000 (Invitrogen) following the manufacturer’s protocols. Cells were harvested 30 hours post-transfection by removing the media, washing once with PBS, and freezing at −80°C. Luciferase assays were performed as before (Kristjansdottir et al. 2015). Firefly luciferase values were first normalized to Renilla values for each well to control for transfection efficiency, and then scaled relative to the normalized firefly values for the control reporter (or to wild-type construct when testing mutant derivatives). Error bars were estimated as the nonparametric equivalent to one standard deviation (NP-SD, ∼68% of the data is within the error bars, e.g. the middle 7 datapoints for n=9).

### Genomic integration using FLP-in system

Flp-In T-REx 293 cells (Life Technologies) were maintained in 10% FBS and 1% penicillin/streptomycin (P/S) supplemented DMEM. Cells were seeded at 750,000 cells/well into 6-well plates using antibiotic free media (DMEM with 10% FBS) and transfected with 250 ng of GFP plasmid and 625 ng of pOG44 (encoding FLP recombinase) using Lipofectamine 2000 (Invitrogen). Media was replaced the following day with DMEM with 10% FBS and 1% P/S (complete DMEM). Selection for integrated cells started 48 hours post transfection by passaging cells into 10 cm plates into complete DMEM with 125 μg/mL hygromycin. Selection continued, with media changes every 3-4 days, until colonies were visible. Colonies were then counted (at least 10 colonies for each cell line) before being passaged into T25 flasks, expanded and frozen.

### Genomic integration of reporters using Bxb1 recombinase

Reporter integration using FLP-in system showed loss of signal over time (data not shown), so we switched to using the Bxb1 recombination system described by the Fowler lab (Matreyek et al. 2017; Matreyek et al. 2020), which is many orders of magnitude more efficient and demonstrated no loss of signal after >3 months of culturing (data not shown). Reporters were integrated into HEK293T cells containing a Bxb1-landing pad, which drives expression from a CMVTetOn promoter. Cells were maintained in 2 µg/mL doxycycline. 250,000 cells were reverse transfected in 6-well plates in DMEM supplemented with 10% FBS, using 3 µg of pCAG-NLS-HA-Bxb1 plasmid (a kind gift from Pawel Pelczar (Addgene plasmid #51271) (Hermann et al. 2014)), 6 µL of Fugene 6 (Promega), and OptiMem (ThermoFisher Scientific) up to 300 µL. 24 hours later, the media was refreshed and cells were forward transfected as before, but instead with 3 µg of the attB-containing plasmid. 24 hours later, the media was changed to DMEM supplemented with 10% FBS and 1% pen-strep. 24 hours later, the media was again changed to DMEM supplemented with 10% FBS, 1% pen-strep and 2 µg/mL doxycycline. Five days after the second transfection, successful integrants (eGFP+/BFP-) were sorted by flow cytometry on a Sony MA900 sorter.

### Flow cytometry

GFP fluorescence of Flp-In T-REx 293 cells expressing integrated acGFP reporters or HEK293T cells expressing integrated eGFP reporters was quantified using at least 100,000 cells with either a BD LSRII flow cytometer using DiVa software (BD Biosciences) or an Attune NxT analyzer, and flow cytometry data was analyzed using FlowJo (TreeStar).

### RNAi knockdown

#### siRNA transfections

HeLa cells were seeded at 75,000 cells/mL in 10 cm culture dishes in DMEM supplemented with 10% FBS, and grown for 24 hours. Cells were then transfected with 30 nM siRNA (UPF1: 5′-GAUGCAGUUCCGCUCCAUUdTdT-3′, 5′AAUGGAGCGGAACUGCAUCdTdT-3′ (Popp and Maquat 2015); SMG6: 5′-CUUGUAAGUAACCUGCAGCUU-3′, 5′-GCUGCAGGUUACUUACAAGUU-3′ (Mascarenhas et al. 2013; Schmidt et al. 2015);

Inert: 5′-UAAAAAUCGCGUGGAUUAAUG-3′, 5′-UUAAUUUACGCGGUUUUUAUU-3′) using the RNAiMAX transfection reagent (Invitrogen) according to manufacturer’s instructions. Transfected cells were seeded the following day on 24-well plates for plasmid transfection for dual luciferase assays (see above).

#### shRNA transductions

Performed as before (Kristjansdottir et al. 2015), except in Flp-In TREx 293 cells expressing integrated yeast-optimized GFP (yGFP) reporters, which were plated at 150,000 cells/mL, and transfected at an MOI (multiplicity of infection) of 10.

### Lentiviral production

500,000 HEK293T cells were plated per well in 6-well plates, using DMEM supplemented with 10% FBS. 24 hours later, cells were transfected with 500 ng psPAX2 (packaging vector), 50 ng VSV-G (envelope vector) and 50 ng of transfer plasmid, using TransIT-LT1 (Mirus) following manufacturer’s instructions. 24 hours later, the media was changed to DMEM supplemented with 1% pen-strep and 20% FBS. 48 hours later, virus was harvested by transferring the conditioned media to a 15 mL Falcon tube, centrifuging at 1000 g for 3 minutes, and then aliquoting the supernatant. Virus was stored at −80°C.

### CRISPRi knockdown

A clonalized dCas9 cell line was generated by transducing 293Ts (containing a Bxb1-integration site (attP) downstream of a CMVTetOn promoter) with lentivirus made from pHR-UCOE-SFFV-dCas9-mCherry-ZIM3-KRAB, a kind gift from Mikko Taipale (Addgene plasmid #154473) (Alerasool et al. 2020). This cell line was single-cell sorted (mCherry+/BFP+) by flow cytometry into 96-well plates, and clones were screened for CRISPRi functionality. Then, lentivirus was produced using the sgRNA plasmids, and 1 mL of lentivirus was used to infect 200,000 cells in a 6-well plate. 48 hours later, 2.5 µg/mL puromycin was added to the media to select for integrants, and the cell lines were maintained in puromycin for at least 5 days before analyzing by flow cytometry.

### RT-qPCR

RNA was isolated from cell culture using Trizol (Thermo Fisher Scientific), following the manufacturer’s instructions. 500 ng of total RNA was used as the template for reverse transcription using RevertAid (Thermo Fisher Scientific), following the manufacturer’s instructions with random nonamers (dN_9_) as the primer, or using a gene-specific primer for Fig S3D. The RT product was diluted 25-fold and 4 µL was used as template in the qPCR reaction, for a total dilution of 50-fold. qPCR reactions were performed using the LightCycler® 480 SYBR Green I Master (Roche), following the manufacturer’s instructions. qPCR reactions were performed in technical triplicates on a LightCycler® 480 Instrument II (Roche). RT-qPCR data was analyzed using the ΔΔCt method, normalized against either GAPDH or PPIA as the housekeeping gene.

### Polysome profiling

For each replicate, four 70% confluent 10 cm plates of cells were treated with 100 μg/mL cycloheximide in media for 3 minutes at 37°C, lifted from the plates in ice cold PBS supplemented with 100 μg/μL cycloheximide, pelleted at 180 *g* for 3 minutes, and lysed in 400 μL ice-cold polysome lysis buffer (10 mM HEPES pH 7.6, 100 mM KCl, 5 mM MgCl_2_, 5 mM DTT, 1% Triton X-100, 100 μg/mL cycloheximide) on ice for 10 minutes. Lysates were clarified by centrifuging at 16,000 *g* for 15 minutes at 4°C and the supernatant was snap frozen in liquid nitrogen. As input, 50 μL of clarified lysate was taken into 1 mL Trizol to represent the cytoplasmic fraction. For polysome profiling, 15-45% (w/v) sucrose gradients in polysome buffer (10 mM HEPES pH 7.6, 100 mM KCl, 5 mM MgCl_2_, 5 mM DTT, 100 μg/mL cycloheximide) were prepared with a Gradient Master (Biocomp). Lysates from five cell lines stably expressing GFP reporters with different length random 3′UTRs were mixed and 500 μL of the mixed lysate was loaded onto the gradient. Lysates were centrifuged at 32,000 rpm for 2.5 hours at 4°C in a SW40 ultracentrifuge rotor. Gradients were then fractionated into equal volume (∼750 μL) fractions with a Brandel density gradient fractionation system and a FoxyJunior fraction collector. From each fraction, 100 μL was taken into 1 mL Trizol for qRT-PCR. For normalization, 0.6 ng of in vitro transcribed Renilla luciferase mRNA was spiked into each fraction. RT-qPCR with 3′UTR-specific primers was then performed.

### Measuring nascent mRNA levels of reporters

Nascent mRNA was isolated using PRO-seq as previously described (Patel et al. 2020), except RT-qPCR (described above) was used to measure mRNA levels of reporters instead of sequencing.

### SILAC-ChIRP-MS

ChIRP-MS was performed as previously described (Chu and Chang 2018), with the following modifications. Twenty 10 cm plates (∼400 million cells) of HEK293 cells per sample were grown in SILAC media for 10 days. SILAC media consists of DMEM for SILAC (ThermoFisher Scientific) supplemented with 10% dialyzed FBS (ThermoFisher Scientific), 1% pen-strep and appropriate SILAC amino acids (ThermoFisher Scientific): L-Lysine-2HCl and L-Arginine-HCl for “light” media, L-Lysine-2HCl (4,4,5,5-D4) and L-Arginine-HCL (^13^C_6_) for “medium” media, L-Lysine-2HCl (^13^C_6_, ^15^N_2_) and L-Arginine-2HCl (^13^C_6_, ^15^N_4_) for “heavy” media. SILAC color per sample was changed between replicates. Each 10 cm plate was then transfected with ∼5 µg (equimolar across plasmids) of GFP reporter plasmid with 400 nt, 2,000 nt, or TRAM1-2,000 nt 3′UTRs using Lipofectamine 3000 reagent (ThermoFisher Scientific) following the manufacturer’s instructions. Transfection provided higher expression of reporters than the integrated eGFP cell lines, which improved ChIRP pulldowns (data not shown). 24 hours after transfection, cells were harvested, pooled and cross linked as described by Chu & Chang (2018). Lysates were sonicated for 4×15 minutes using a Bioruptor UCD-200 (Diagenode). Hybridization of probes was performed for 16 hours. Otherwise all pulldown steps were performed as described (Chu and Chang 2018). Mass spectrometry was performed by the Proteomics Facility in the Biotechnology Resource Center at Cornell University as follows:

#### In-solution trypsin digestion of SILAC labeled samples

In solution digestion was performed on S-Trap micro spin column (ProtiFi, Huntington, NY, USA) following an Strap protocol on as described previously (Yang et al. 2018).

#### Nano LC/MS/MS Analysis on Orbitrap Fusion

The SILAC tryptic digests were reconstituted in 50 μL of 0.5% FA estimated at 0.1 µg/µL for nanoLC-ESI-MS/MS analysis, which was carried out using an Orbitrap FusionTM TribridTM (Thermo-Fisher Scientific, San Jose, CA) mass spectrometer equipped with a nanospray Flex Ion Source, and coupled with a Dionex UltiMate3000RSLCnano system (Thermo, Sunnyvale, CA) (Harman et al. 2018; Yang et al. 2018). The tryptic peptide samples (5 μL) were injected onto a PepMap C-18 RP nano trap column (5 µm, 100 µm i.d x 20 mm, Dionex) with nanoViper Fittings at 20 µL/min flow rate for on-line desalting. The peptides were separated on a PepMap C-18 RP nano column (2 µm, 75 µm x 25 cm) at 35°C, in a 120 min gradient of 5% to 38% acetonitrile (ACN) in 0.1% formic acid at 300 nL/min and followed by a 8 min ramping to 90% ACN-0.1% FA and a 9 min hold at 90% ACN-0.1% FA. The column was re-equilibrated with 0.1% FA for 25 min prior to the next run. The Orbitrap Fusion is operated in positive ion mode with spray voltage set at 1.6 kV and source temperature at 275°C. External calibration for FT, IT and quadrupole mass analyzers was performed. In data-dependent acquisition (DDA) analysis, the instrument was operated using FT mass analyzer in MS scan to select precursor ions followed by 3 second “Top Speed” data-dependent CID ion trap MS/MS scans at 1.6 m/z quadrupole isolation for precursor peptides with multiple charged ions above a threshold ion count of 10,000 and normalized collision energy of 30%. MS survey scans at a resolving power of 120,000 (fwhm at m/z 200), for the mass range of m/z 375-1575. Dynamic exclusion parameters were set at 50 s of exclusion duration with ±10 ppm exclusion mass width. All data were acquired under Xcalibur 3.0 operation software (Thermo-Fisher Scientific).

#### Mass spectrometry data analysis

All MS and MS/MS raw spectra were processed and searched using Sequest HT software within the Proteome Discoverer 2.4 (PD 2.4, Thermo Scientific). The Homo sapiens NCBI database containing 81,725 entries was used for database searches. The database search was performed under a search workflow with the “Precursor Ions Quantifier” node for SILAC 3plex (R0K0, R6K4, R10K8) quantitation. The default setting for protein identification in Sequest node were: two mis-cleavages for full trypsin with fixed carbamidomethyl modification of cysteine, variable modifications of 6.020 and 10.008 Da on Arginine, 4.025 and 8.014 Da on lysine, N-terminal acetylation, methionine oxidation and deamidation of asparagine and glutamine residues. The peptide mass tolerance and fragment mass tolerance values were 10 ppm and 0.6 Da, respectively. Only high confidence peptides defined by Sequest HT with a 1% FDR by Percolator were considered for the peptide identification. The SILAC 3-plex quantification method within PD 2.4 was used with unique + razor peptides only to calculate the heavy/light ratios using pairwise ratio based for all identified proteins without normalization and exclude the deamidation modification. The final protein group list was further filtered with two peptides per protein in which only #1-ranked peptides within top scored proteins were used.

### Half-life measurements

To create a Bxb1 landing pad cell line with a repressible promoter, HEK293T cells were transduced at low MOI with lentivirus consisting of (1) a repressible TRE promoter (Tet-Off), followed by a Bxb1 attP site and a BFP marker, and (2) a CMV promoter driving constitutive expression of the doxycycline-responsive tTa transactivator. To generate a clonal cell line, BFP+ cells were single-cell sorted by FACS into 96-well plates. Candidate clones were screened for BFP expression, efficient repression by doxycycline, and high recombination efficiency, and a single clone was chosen for subsequent decay assays. eGFP reporters with different length 3′UTRs were then integrated into this cell line using Bxb1 recombinase (described above).

Five cell lines expressing eGFP with different 3′UTRs under the control of a doxycycline-repressible promoter were used to measure reporter half-lives. Cell lines were seeded in triplicate (per timepoint) in 6-well plates, with 200,000 cells per well. First, reporter transcription was shut off with a 48 hour 1 ng/µL doxycycline treatment. Then, a pulse of transcription was induced by removing doxycycline from the media for 4 hours. Finally, transcription was shut off using 2 µg/mL doxycycline, and time points were taken at 0, 0.5, 1, 2, 3, 4, 8 and 12 hours post-addition of doxycycline. At the appropriate timepoint, 1 mL of Trizol was added per well, and RNA was isolated for each cell line expressing a different 3′UTR reporter. RNA isolation and RT-qPCR were performed as described above. Ct values were normalized to PPIA, and then to the 0 hour time point for each cell line expressing a different 3′UTR reporter. Half-life measurements were performed in biological triplicates, and qPCR was performed in technical triplicate.

### RNA FISH

Stellaris® FISH Probes recognizing either the coding region of eGFP mRNA or GAPDH mRNA labeled with Quasar® 670 Dye (VSMF-1015-5 for eGFP and SMF-2019-1 for GAPDH, from Biosearch Technologies, Inc., Petaluma, CA) were hybridized to HEK293T cell lines expressing integrated eGFP reporters with different length 3′UTRs according to manufacturer’s protocol. Briefly, cells were seeded onto cover slips pretreated with poly-D-lysine and cultured overnight in DMEM supplemented with 10%FBS and 1%P/S at 37°C and 5% CO_2_. Cells were washed with PBS and fixed with 3.7% formaldehyde in PBS for 10 min at room temperature. Fixed cells were washed twice with PBS. Cells were permeabilized in 70% ethanol for at least 1 hour at 2-8°C. Cells were subsequently washed with Wash buffer A (10% formamide in 1X Stellaris RNA FISH Wash Buffer A) before staining with the probes in hybridization buffer (2 µL of probes/cover slip, 10% formamide in Stellaris RNA FISH Hybridization Buffer) overnight in a humidified chamber at 37°C in the dark. After incubation, cells were washed with Wash Buffer A and stained with DAPI nuclear stain for 30 min in the dark at room temperature. Cover slips were subsequently washed with Wash buffer B for 5 minutes and mounted onto the slides with Vectashield® Mounting Medium (H-1000, Vector Laboratories). Images were taken at the Imaging Facility, Biotechnology Resource Center, Cornell University using a Nikon Inverted microscopy. Images were taken at 20 or 40X with fixed settings for DAPI, FITC, and Alexa 647 channels, with single slices through the z-axis. RNA FISH signal was analyzed using ImageJ, and nuclear/cytoplasmic ratios were calculated as previously described (Kelley and Paschal 2019).

## Acknowledgements

We thank the Proteomics Facility of Cornell University (CU) (RRID:SCR_021743) for providing the mass spectrometry data and NIH SIG grant 1S10 OD017992-01 grant support for the Orbitrap Fusion mass spectrometer. We also thank the Flow Cytometry Facility of CU (RRID:SCR_021740), Imaging Facility of CU (RRID:SCR_021741), and the Genomics Facility of CU (RRID:SCR_021727) for their support. C.W.P.D was supported by a fellowship from Boehringer Ingelheim Fonds.

## Author Contributions

C.W.P.D., K.K., and A.G. conceived and planned the experiments. C.W.P.D., K.K., J.D.W., L.T.V., E.A.F., and R.G. carried out the experiments. C.M. and A.K. provided technical support. C.W.P.D. and K.K. performed data analysis. C.W.P.D., K.K., and A.G. interpreted the results and wrote the manuscript. All authors provided critical feedback and helped shape the research.

---

**Fig S1.**
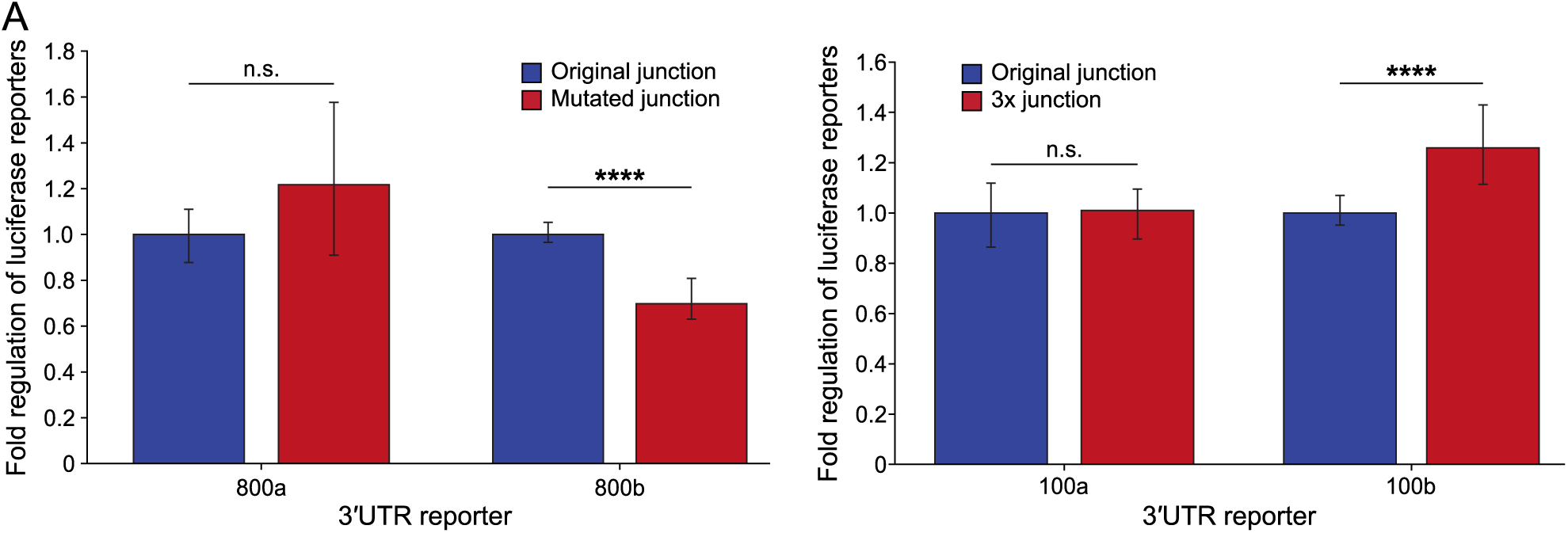
Cloning junction does not repress reporters with random 3′UTRs. A) Luciferase assays of indicated random 3′UTR reporters in A549 cells, normalized to the reporter with the original junction (n=9, error bars are NP-SD). (Left) The effect of mutating the 6 nt cloning junctions on two different 800 nt random 3′UTR reporters. (Right) The effect of adding three copies of the 6 nt cloning junction on two different 100 nt random 3′UTR reporters. ****p<0.0001; p-values derived from two-tailed t-tests.

**Fig S2.**
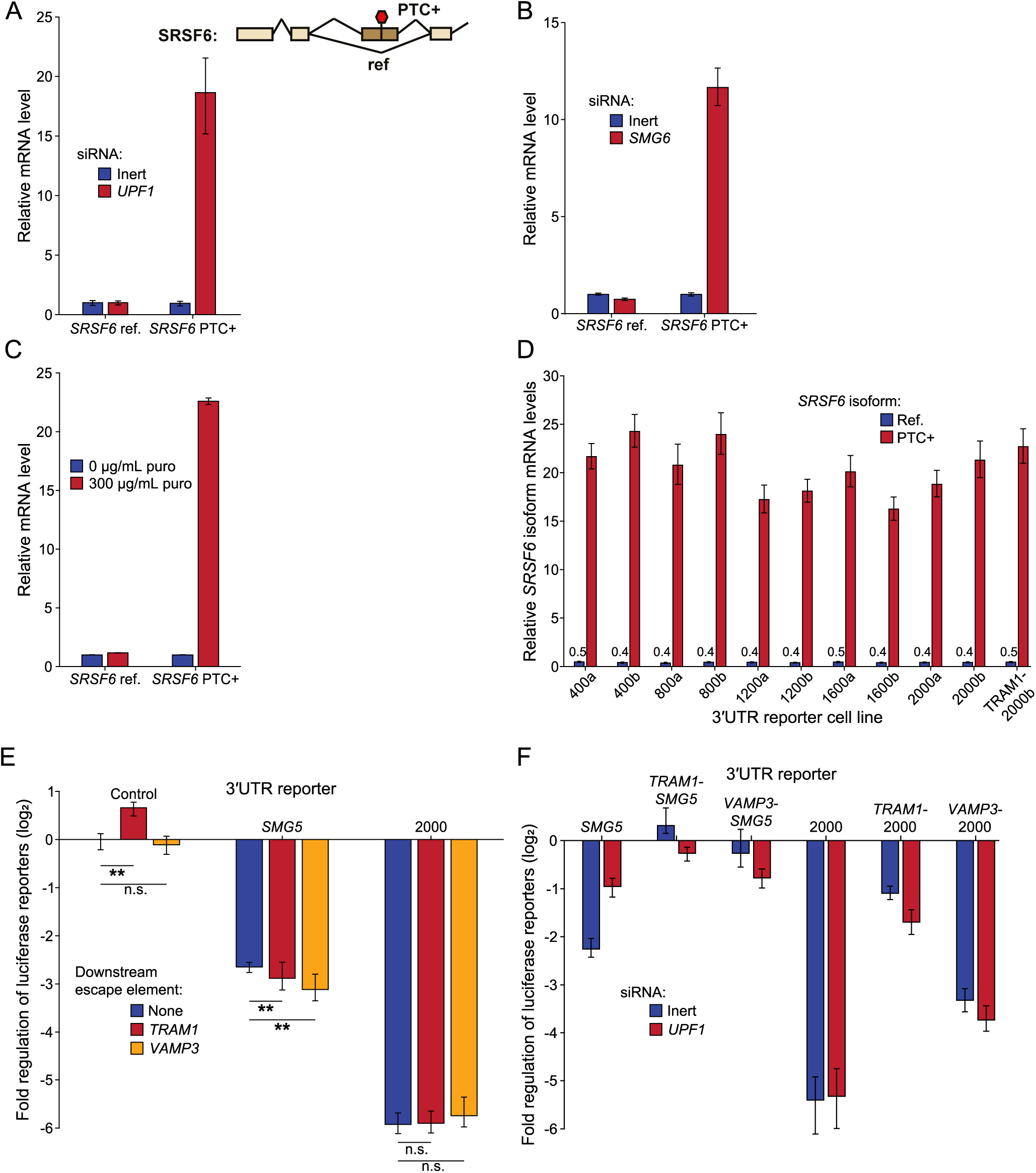
NMD does not cause repression of long 3′UTR reporters; validation of NMD inhibition. A) (Top) Schematic showing alternative splicing isoforms of *SRSF6*. The “PTC+” isoform contains a premature termination codon in the third retained exon, whereas the “ref” isoform does not contain a PTC due to exon skipping. (Bottom) Change in mRNA levels for the two *SRSF6* isoforms after 72 hour siRNA knockdown of *UPF1*, as measured by RT-qPCR (n=9, two independent siRNA transfections, each measured in triplicate, error bars indicate one SD). Each isoform levels are normalized to the inert siRNA treatment. B) As in (A), but after siRNA knockdown of *SMG6*. C) As in (A), but after 8 hour treatment with 300 µg/mL puromycin. D) As in (A), but after 4 hour treatment with 0.5 µM SMG1i in HEK293T cells. *SRSF6* isoform levels shown for each cell line expressing a different length integrated random 3′UTR eGFP reporter, normalized to DMSO vehicle treated control (three biological replicates, each in technical triplicate, error bars indicate one SD). E) As in Figure 2D, but escape elements were instead cloned into the 3′ end of the reporter 3′UTRs (n=6, error bars are one NP-SD). F) The effect of *UPF1* knockdown on expression of transiently transfected luciferase reporters with *SMG5* or 2,000 nt random 3′UTRs, with or without upstream *TRAM1* or *VAMP3* escape elements. Reporters are normalized to the control 100 nt 3′UTR reporter with no escape element (n=9, two independent siRNA transfections, error bars are NP-SD). **p<0.01; p-values derived from two-tailed t-tests.

**Fig S3.**
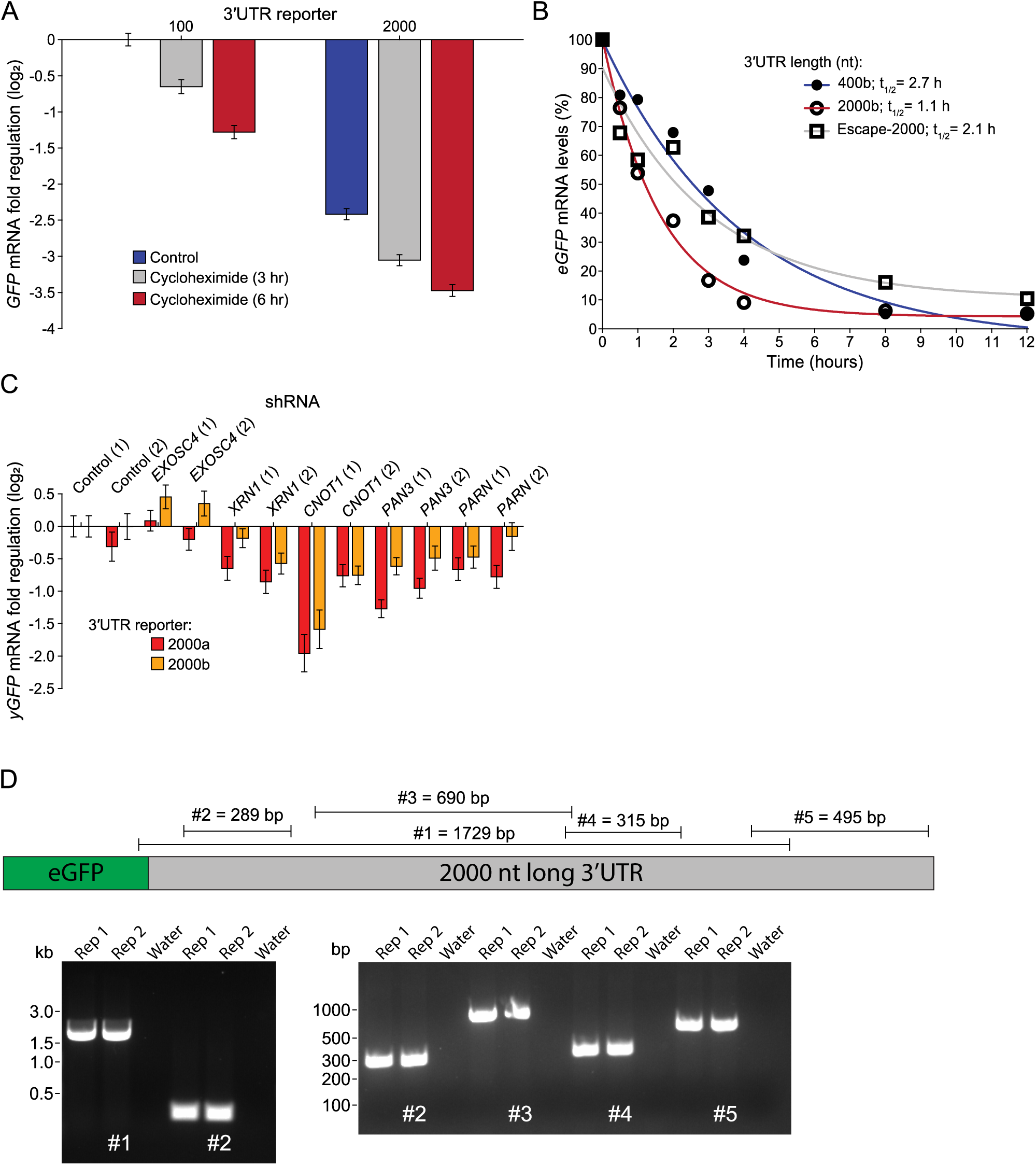
LMD changes half-life of long 3′UTR reporters, but does not involve major deadenylases and exonucleases. A) The effect of translation inhibition by a 3 or 6 hour treatment with cycloheximide on the mRNA levels of two integrated yGFP reporters with different 2,000 nt random 3′UTRs in T-REx HEK293FLP cells, relative to a control 100 nt 3′UTR reporter (three biological replicates each in technical triplicate, error bars are one SD). B) mRNA decay curves of integrated eGFP reporters with different length random 3′UTRs expressed from the doxycycline repressible Tet-OFF promoter in HEK293T cells, measured by RT-qPCR (three biological replicates each in technical triplicate). Half-lives of each reporter noted in legend. The Escape-2000 reporter is the same as in Fig 3F. C) The effect of exonuclease or deadenylase knockdown with shRNAs on the mRNA levels of integrated yGFP reporters with two different 2,000 nt random 3′UTRs, normalized first to a 100 nt control 3′UTR then to the respective 2000 nt random 3′UTR reporter treated with Control (1) shRNA (two biological replicates each in technical triplicate, error bars represent one SD). D) (Top) Diagram showing binding locations of primer pairs across the 2000 nt random 3′UTR used in ChIRP-MS. (Bottom) Agarose gel results of RT-PCR using primers binding along the breadth of the random 2000 nt long 3′UTR used in ChIRP-MS, showing no part of the 3′UTR is removed through splicing.

**Fig S4.**
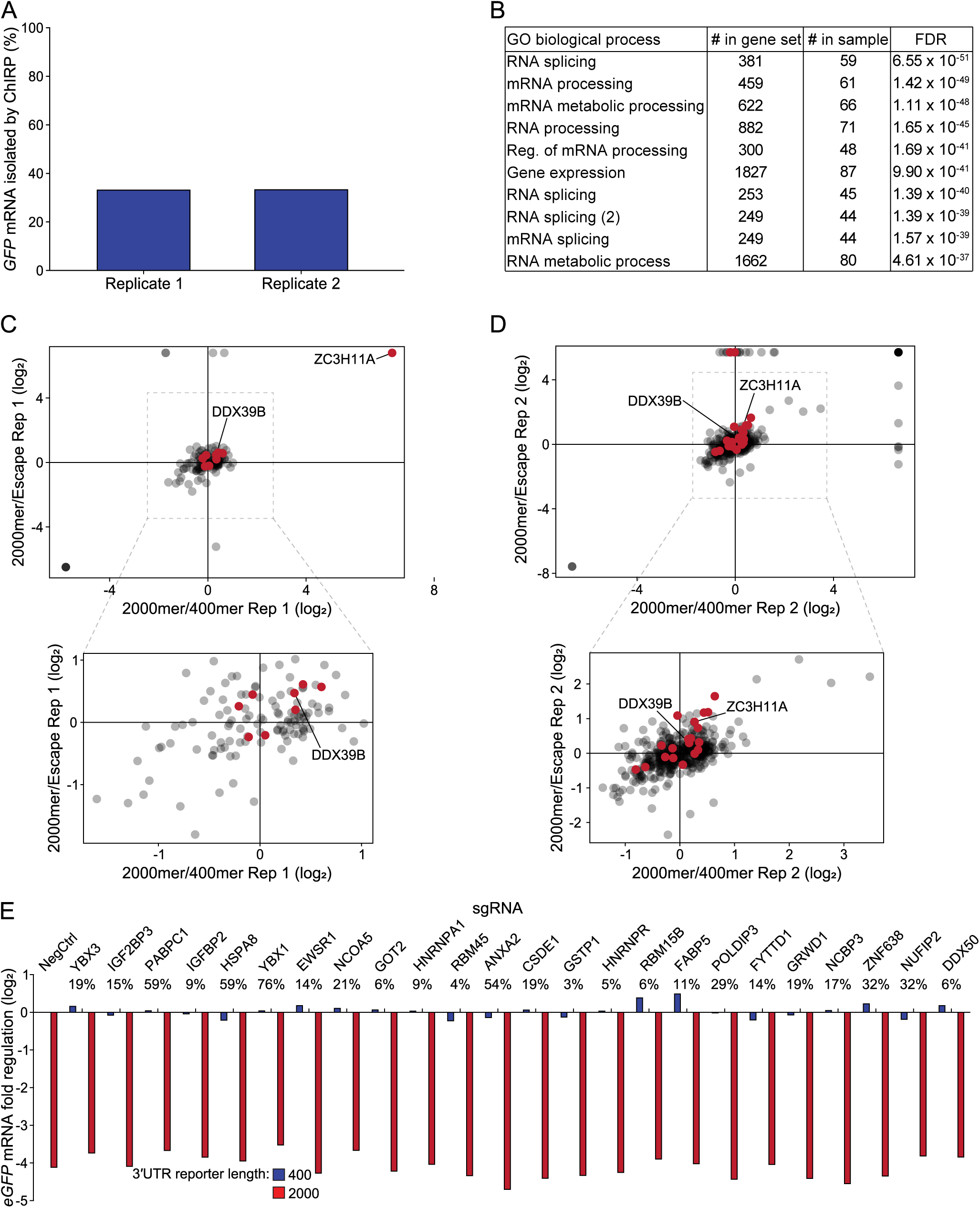
Screening LMD candidate factors identified by SILAC-ChIRP-MS. A) Percent of *GFP* mRNA recovered by SILAC-ChIRP-MS across two replicates with different SILAC labelling strategies, relative to the starting amount before pulldown, measured by RT-qPCR (one biological replicate in technical triplicate). B) Top ten terms from gene ontology (GO) analysis of proteins identified by mass spectrometry across all three samples, for one SILAC-ChIRP-MS replicate. C) SILAC abundance ratios of proteins associated with a GFP reporter with a 2,000 nt random 3′UTR relative to a 400 nt random 3′UTR (x axis), compared to the proteins associated with a GFP reporter with a 2,000 nt random 3′UTR relative to a 2,000 nt random 3′UTR with an upstream *TRAM1* escape element (y axis), for a single SILAC-ChIRP-MS replicate. Bottom shows zoomed view of boxed area. Red dots indicate proteins part of the gene ontology term “mRNA export from nucleus”. D) As in (C), but for the second SILAC-ChIRP-MS replicate. E) LMD candidate factors were chosen from SILAC-ChIRP-MS data shown to be strongly enriched or depleted on a GFP reporter with a 2,000 nt random 3′UTR compared to reporters with either a 400 nt random 3′UTR or a 2,000 nt random 3′UTR with an upstream *TRAM1* escape element. The effect of candidate factor knockdowns, by CRISPRi, on LMD was tested by measuring mRNA level changes in expression of integrated eGFP reporters with a 400 nt or 2,000 nt random 3′UTR by RT-qPCR (one biological replicate, in technical triplicate). Here, an example subset of the 70 total screened factors is shown. All samples are normalized to the cell line expressing a 400 nt random 3′UTR and the negative control sgRNA (NegCtrl). Percentages show the mRNA levels of each candidate factor after knockdown, relative to the cell line expressing a negative control sgRNA.

**Fig S5.**
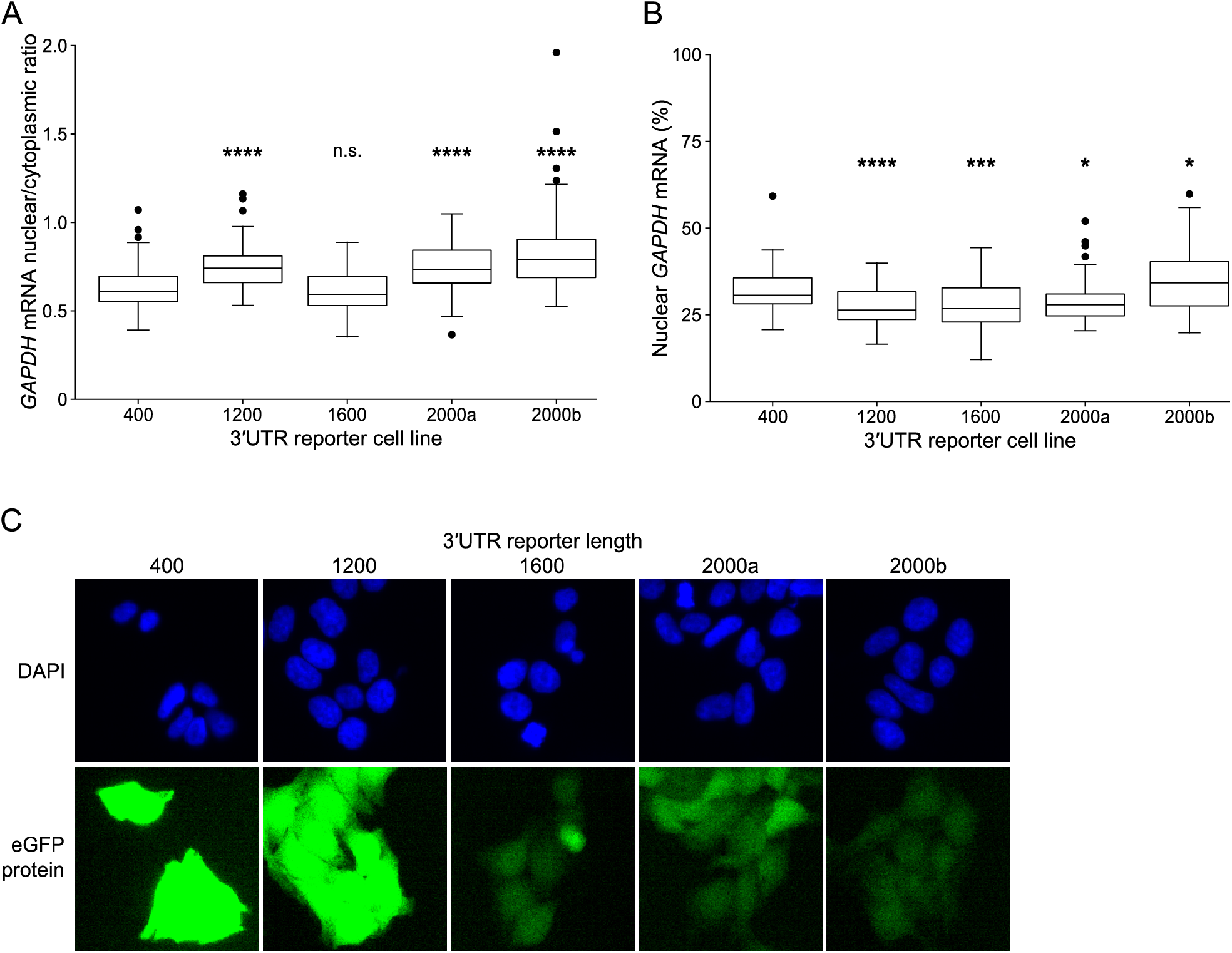
Controls for RNA FISH data. A) Nuclear/cytoplasmic ratios of *GAPDH* mRNA in HEK293T cell lines expressing integrated eGFP reporters with different length random 3′UTRs, as measured by RNA FISH (n≥50 cells). Statistical tests compare the cell line expressing 400 nt long 3′UTR reporter against indicated cell line. B) The percent of *GAPDH* mRNA localized to the nucleus in HEK293T cell lines expressing integrated eGFP reporters with different length random 3′UTRs, as measured by RNA FISH (n≥50 cells). Statistical tests compare the cell line expressing 400 nt long 3′UTR reporter against indicated cell line. C) Representative images (same field of view as in Fig 5C) of DAPI stained (top) and eGFP fluorescing (bottom) HEK293T cells expressing integrated eGFP reporters with different length random 3′UTRs. *p<0.05, ***p<0.001, ****p<0.0001; p-values derived from two-tailed t-tests.

